# REACTIVATION PROTECTS MOTOR MEMORIES FROM INTERFERENCE BY COMPETING LEARNING

**DOI:** 10.1101/2025.06.26.661747

**Authors:** Tharan Suresh, Adarsh Kumar, Pratik Mutha

## Abstract

Memories are often thought to become increasingly stable and resistant to change over time. The reconsolidation framework proposes that reactivating a stable memory transiently increases its susceptibility to modification by subsequent experience, but evidence for such reactivation-induced destabilization remains mixed. Here we tested whether reactivation alters the susceptibility of a visuomotor memory to interference from competing learning. Across a series of motor adaptation experiments, we briefly reactivated a previously acquired memory before introducing an opposing perturbation. Contrary to reconsolidation predictions, reactivation did not increase interference during later retrieval. Instead, reactivation reduced interference when the competing memory was not allowed to stabilize after acquisition. Our findings challenge the view that reactivation universally destabilizes motor memories and instead suggest that it shapes which competing motor memory is expressed at retrieval.

## INTRODUCTION

A central question in neuroscience is how new learning interacts with and modifies previously acquired memories. Across many systems, memories are thought to stabilize over time (Brashers-Krug et al., 1996; Dudai, 2004; Dudai et al., 2015; McGaugh, 2000), become distributed over different neural circuits (Doyon & Benali, 2005; Hikosaka et al., 1999, 2002; Nudo et al., 1996) and grow increasingly resistant to modification from new learning that might compete for the same neural resources (Dudai, 2004; Lee, 2009; Nader et al., 2000a). This notion of relative permanence however has been challenged by a line of work demonstrating that an initially acquired memory can indeed be altered (Dudai, 2006; Elsey et al., 2018; Nader & Hardt, 2009; Sara, 2000; Schiller, 2022). The “reconsolidation” framework proposes that a previously acquired memory can be brought to a temporarily labile state through reactivation or active retrieval, potentially increasing its vulnerability to change via subsequent learning (Dudai & Eisenberg, 2004; Schlichting & Preston, 2014). Early support for this idea came from rodent fear conditioning studies showing that a consolidated memory can be corrupted if neural processing is disrupted after reactivation (Nader et al., 2000b), a finding that has been reported in humans as well (Kindt et al., 2009). Subsequent work has reported reactivation-mediated memory modification in object recognition memory in rodents (Davis et al., 2010; Winters et al., 2011), as well as declarative memory (Chan & LaPaglia, 2013) and sequence learning in humans (Gabitov et al., 2017, 2019; Walker et al., 2003). Notably though, there is growing evidence that such reactivation-induced modification is not always observed (Censor et al., 2016; de Beukelaar et al., 2014; Hardwicke et al., 2016; Potts & Shanks, 2012; for a review, see Schroyens et al., 2023), which has led to a major debate about the generality of reconsolidation effects across memory systems, and more fundamentally, about whether behavioral signatures such as interference necessarily reflect changes in memory stability.

Sensorimotor adaptation provides a powerful system to address this issue because interactions between competing motor memories can be directly examined using interference paradigms. In such studies, participants first learn to compensate for a perturbation (task A) and are subsequently exposed to an opposing perturbation (task B). The learning of task B disrupts the expression of the previous A memory when it is later tested, demonstrating behavioral interference. Classic studies employing this approach have shown that interference can occur even if the first memory is acquired up to a week before new learning (Caithness et al., 2004; Krakauer et al., 2005; Krakauer, 2009; Kumar et al., 2018; Miall et al., 2004). Importantly, this interference arises *without* explicit reactivation of the original memory prior to the acquisition of B. This suggests that interference in such paradigms need not necessarily arise from destabilization of a previously stored memory, but may reflect competition between multiple motor memories at retrieval rather than changes to the underlying memory trace. This leaves open the question of whether reactivation alters the underlying stability of the original memory, or whether it modulates how competing memories are expressed at retrieval.

We therefore asked how reactivation influences interference between human motor memories. We tested the hypothesis that reactivation of a previously acquired memory trace renders it temporarily labile, thereby increasing its susceptibility to degradation by subsequent learning. Across a series of motor adaptation experiments, we briefly reactivated a stable motor memory before exposing participants to an opposing perturbation. Contrary to classic reconsolidation predictions, reactivation did not increase interference. Instead, reactivation enhanced resistance to interference when the competing memory was prevented from stabilizing, indicating that rather than uniformly promoting destabilization, its effects depend on how subsequent learning is structured. These results thus refine current theories of reconsolidation and suggest that reactivation influences which competing memory is expressed at retrieval.

## METHODS

### Participants

We recruited 140 young, right-handed healthy adults (80 males, 60 females, aged 18 to 40 years) across three experiments. Out of these, 60 subjects were randomly assigned to 3 groups in experiment 1 (n = 20 each), 40 were randomly distributed across 2 groups in experiment 2 (n = 20 each), and the remaining 40 were randomly allocated to 2 groups in experiment 3 (n = 20 each). The sample size of 20 subjects per group was arrived at based on past work (Kumar & Mutha, 2023; Song & Bédard, 2015) that found reliable adaptation (Cohen’s *d* > 1 or *η*^2^_*p*_ > 0.25) with much smaller group sizes (n <= 13). Handedness was determined by the Edinburgh Handedness Inventory (Oldfield, 1971). Participants had no self-reported history of neurological or psychiatric illness, brain injury, or addiction. Experiments were conducted after obtaining written informed consent, and the protocol was approved by the institutional ethics committee.

### Experimental Setup and Task

We used a virtual reality setup where participants sat facing a large digitizing tablet (GTCO CalComp) on which they made planar reaching movements from a circular start position (green color, 1.5 cm diameter) to target circles (red color, 1.5 cm diameter) using a hand-held stylus (Figure 1a). A high-definition television screen, positioned above the tablet and facing downwards, was used for presenting the start and target positions. To eliminate direct visual feedback of the hand, a semi-silvered mirror was placed between the television and the digitizing tablet. Feedback about hand position was instead made available via a cursor displayed on the screen along with the start position and targets. This cursor feedback could be aligned with the actual hand position or rotated relative to the direction of hand motion (visuomotor rotation).

**Figure 1.**
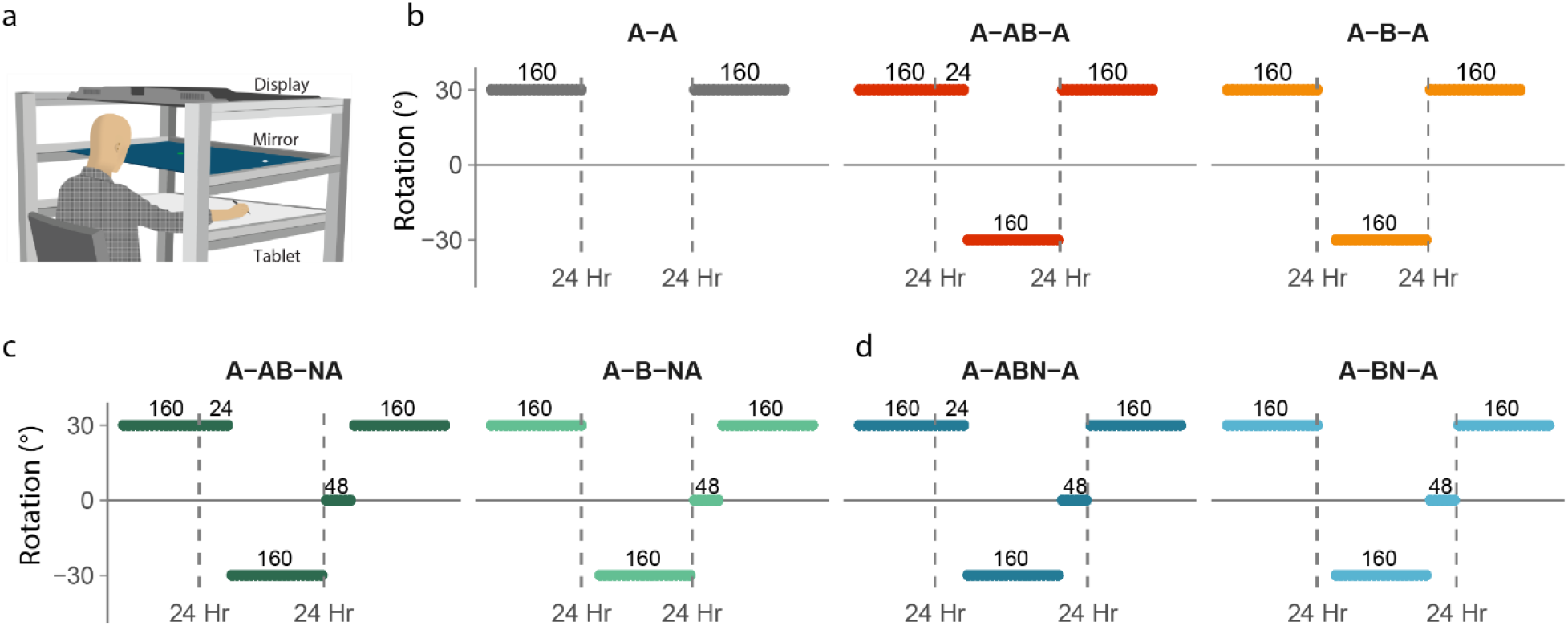
**(a)** Experimental setup comprising of a virtual reality system that restricted movements to the horizontal plane. Participants performed arm-reaching movements on a digitizing tablet while looking into a mirror placed between the tablet and a horizontally mounted high-definition display. Feedback about hand position was displayed via the display onto the mirror using a cursor. **(b-d)** Trial structure for the different groups of (b) experiment 1, (c) experiment 2 and (d) experiment 3. Across experiments participants performed variants of a 3-day A-B-A visuomotor adaptation task, wherein following baseline practice, they learned a 30° counterclockwise rotation A on day 1, followed by a competing 30° clockwise rotation B on day 2, and were retested on A on day 3, either with or without reactivation.

Participants performed 10 cm long reaching movements from the start position to the target circle. Each trial began by participants bringing the stylus into the start circle and staying there for 250 ms, after which one of eight radially arranged targets (equally spaced around an imaginary ring of 10 cm radius) appeared simultaneously with an audio “go” cue. The order of target presentation was pseudo-randomized such that a target appeared only once per cycle of 8 consecutive trials. Participants were asked to move as quickly and accurately as possible to the displayed target. Each trial lasted for a maximum of 2 sec, and if the participant exceeded the time limit, that trial was ended. A score based on the spatial accuracy of the reaching movement was displayed on the screen at the end of each trial. Participants received 10 points for accurately bringing the stylus into the target, 5 points if their reach ended outside the target but within a 2.5 cm radius of the target’s center, and no points for reaches that ended beyond a 2.5 cm radius from the target center. Points were used to maintain participant motivation; they were not analyzed and did not influence the compensation participants received.

We used a three-day “A-B-A” paradigm that involved learning a 30° counterclockwise rotation A on day 1, followed by adaptation to a competing 30° clockwise rotation B with or without reactivation of the previously consolidated learning of A on day 2, and retesting on A on day 3. Participants started each learning phase at roughly similar times of the day to minimize the effects of circadian rhythm and homeostatic factors. Before learning A on day 1, participants performed 48 no-perturbation “baseline” trials to familiarize themselves with the setup and the task.

#### Experiment 1

Participants in experiment 1 were randomly assigned to one of three groups. Participants in the A-B-A group learned rotation A over a block of 20 cycles (1 cycle = 8 trials) on day 1, rotation B over 20 cycles on day 2 and were retested on rotation A over 20 cycles on day 3 (Figure 1b). In contrast, participants in the A-AB-A group learned A on day 1, practiced A again for 3 cycles on day 2 to briefly reactivate the previously acquired memory and then learned B immediately thereafter. This was followed by retesting on A on day 3. Participants in the A-A group learned A on day 1 and were retested on A over 20 cycles on day 3 like the other two groups; this group did not learn B on day 2. Each participant made 20 movements to each target during training.

#### Experiment 2

In experiment 2, we introduced null rotation (N) trials prior to A relearning on day 3 in two new groups. Participants in the A-B-NA group learned A on day 1 (20 cycles), B on day 2 (20 cycles), and were retested on A on day 3 (20 cycles) after performing 6 “washout” cycles (Figure 1c). No rotation was applied on the washout trials. Washout was done to reduce any residual adaptation of B to near baseline levels and to remove potential anterograde effects from it on subsequent A learning (Caithness, 2004; Krakauer et al., 2005). In contrast, participants in the A-AB-NA learned A on day 1 over 20 cycles, reactivated A (3 cycles) and then immediately learned B (20 cycles) on day 2, and were finally retested on A (20 cycles) on day 3 following 6 cycles of washout trials.

#### Experiment 3

Our third experiment was like experiment 2, except that we introduced the null rotation washout trials immediately after B learning on day 2. Thus, participants in the A-BN-A group learned A on day 1 (20 cycles), learned B (20 cycles) followed by 6 washout cycles on day 2, and were retested on A (20 cycles) on day 3 (Figure 1d). In contrast, participants in the A-ABN-A first learned A (20 cycles) on day 1. On day 2, they reactivated A over 3 cycles, then learned B (20 cycles) and then performed 6 cycles of washout trials. This was followed by retesting on A (20 cycles) on day 3.

### Data Analysis

We recorded the hand (stylus) and cursor positions for offline analysis. Kinematic data were filtered using a low-pass Butterworth filter with a cutoff frequency of 10 Hz. Velocity was calculated by differentiating position data. Adaptation to the visuomotor rotation was quantified as a reduction of direction error across learning cycles. Direction error was defined as the angle between the line connecting the start and target circles and the line connecting the start circle and hand position at peak velocity. Direction errors were baseline corrected by subtracting the mean direction error on the last two familiarization cycles from all trials. We focused on the direction errors during the early and late phases of learning, which were defined as the mean direction error on the first and last learning cycle respectively.

Across all three experiments, nine participants (no more than two participants in any single group) had to be excluded from the analysis for protocol violations, including a failure to follow experimenter instructions resulting in highly aberrant movements during A learning on day 1. For the remaining 131 participants, trials where they did not move, or failed to start a trial within the stipulated time, or lifted the stylus off the tablet leading to a loss of data, were labeled as “bad trials” and excluded. In addition, trials where direction errors were more than 3 standard deviations away from the mean direction error computed for each learning phase within each group were treated as outliers and excluded. Across all participants included in the final analysis (n = 58 for experiment 1, 37 for experiment 2 and 36 for experiment 3), a total of 1.3% of trials were removed based on all the above criteria combined. Statistical comparisons were done using mixed ANOVAs, followed by Holm-Bonferroni-corrected post-hoc tests when appropriate. Details of specific comparisons are given along with the corresponding results. The significance threshold for all comparisons was set at 0.05.

## RESULTS

### Interference was not increased by reactivation

We first tested whether reactivation increases the susceptibility of a previously learned motor memory to interference from competing learning. In experiment 1, participants were divided into three groups: A-AB-A (n = 19) who briefly reactivated A on day 2 before learning B, A-B-A (n = 19) who learned B without reactivation, and A-A (n = 20) who did not learn B and served as a control. All groups relearned A on day 3.

Figure 2a shows the learning curves for all groups across all 3 days of the experiment. Day 1 learning of rotation A was very similar across groups, with direction error decreasing from approximately 27° early on to about 8° by the end of the learning phase. We did not find any statistical differences in A learning across groups (Figure 2b). Our group x time ANOVA revealed a significant effect of time (F_(1,55)_ = 881.72, p < 0.001, ω^2^ = 0.82), but no effect of group (F_(2,55)_ = 0.36, p = 0.697, ω^2^ = 0) or interaction (F_(2,55)_ = 0.27, p = 0.763, ω^2^ = 0). Twenty-four hours after the initial learning of A, participants in the A-AB-A group actively practiced A again for a brief period of 24 trials (3 cycles). Direction errors were lower (Figure 2c) from the outset of the reactivation period (14.269±1.136°) compared to early naïve A learning on day 1 (27.324±0.705°). This indicated that the previously acquired A memory had been retained and could be retrieved. By the end of the reactivation period, errors were comparable to those seen at the end of A learning on day 1. This was supported by a significant day (day 1, day 2) x time (early, late) interaction (F_(1,18)_ = 38.434, p < 0.001, ω^2^ = 0.363), with post-hoc tests showing smaller direction errors at the early stage (t_(18)_ = 9.632, p < 0.001, CI for mean difference = [9.039, 17.069]) but not at the end (t_(18)_ = 1.210, p = 0.242, CI for mean difference = [-2.089, 4.976]) of naïve learning (day 1) and reactivation (day 2) periods.

**Figure 2.**
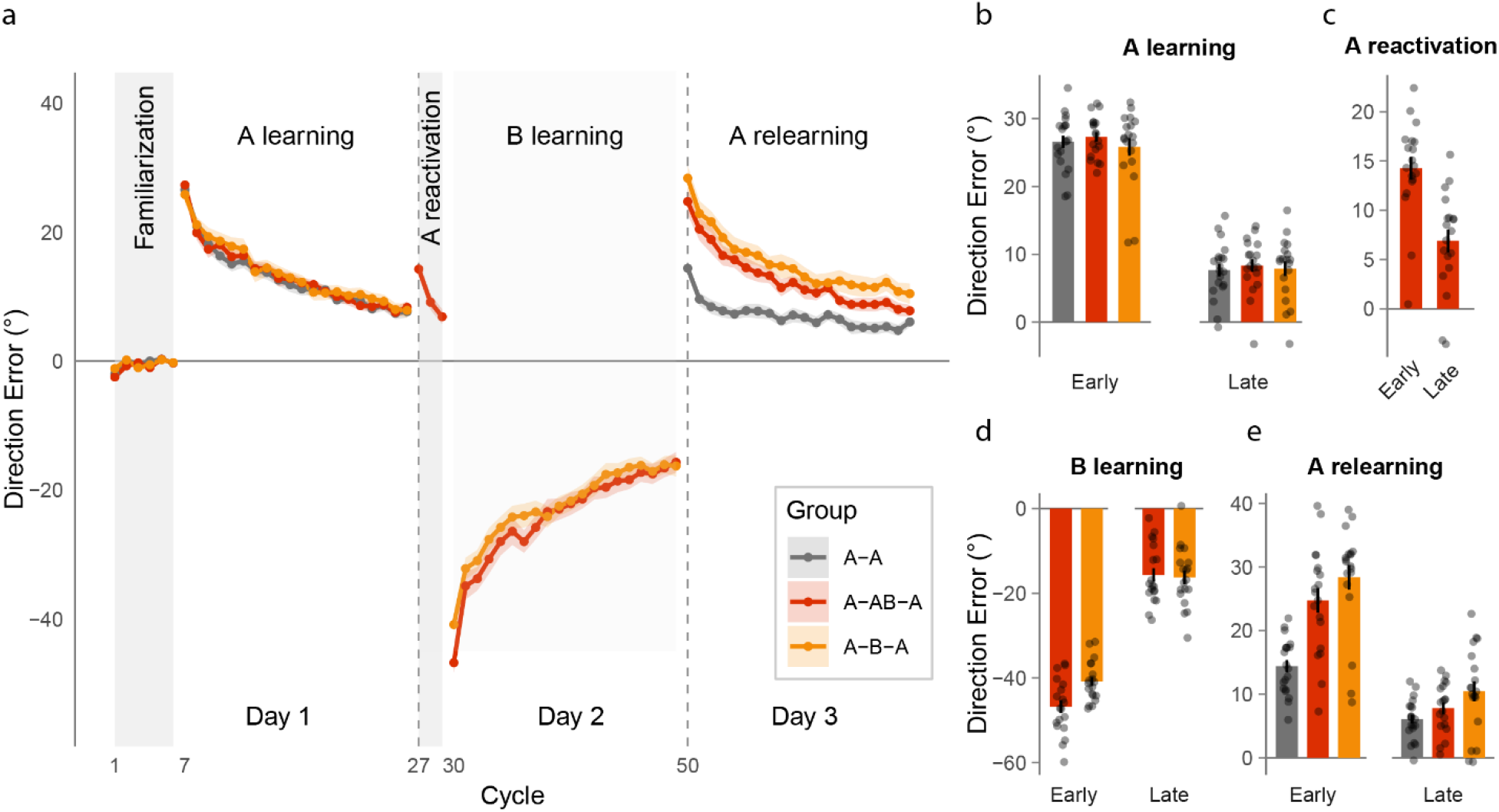
**(a)** Learning curves showing mean direction error as a function of cycles, plotted for the different blocks of experiment 1. The reactivation group (A-AB-A) is shown in red, the no-reactivation group (A-B-A) in orange, and the control group (A-A) in gray. Dark vertical dashed lines represent 24-hour intervals between the blocks. Shaded areas represent SEM. **(b, c, d, e)** Mean ± SEM direction error during early (first 8 trials) and late learning (last 8 trials) phases for each learning block. Dots represent individual participants.

Following reactivation, participants in the A-AB-A group were exposed to the opposing rotation B. This group showed aftereffects of practicing A on day 2 (Figure 2d), which resulted in larger direction errors during the early B learning phase (−46.766±1.501°) than the A-B-A group that did not reactivate (−40.854±1.085°). Errors decreased across trials in both groups, and no differences were evident at the end of learning (Figure 2d). This was supported by a significant group x time interaction (F_(1,36)_ = 5.057, p = 0.031, ω^2^ = 0.050), with post-hoc tests showing significant differences in direction errors during early learning (t_(36)_ = -3.192, p = 0.006, CI for mean difference = [-11.083, -0.742]), but not at the end (t_(36)_ = 0.250, p = 0.804, CI for mean difference = [-5.767, 6.903]).

We next asked whether reactivation influenced relearning of A on day 3. When comparing A learning across groups on day 3, we noted a significant group x time interaction (F_(2,55)_ = 13.149, p < 0.001, ω^2^ = 0.085). Post-hoc tests indicated significantly smaller direction errors for the A-A group relative to the A-B-A group (t_(55)_ = -5.987, p < 0.001, CI for mean difference = [-21.138, - 6.812]) as well as the A-AB-A group (t_(55)_ = -4.433, p < 0.001, CI for mean difference = [-17.510, -3.184]) during early learning (Figure 2e). Critically, relearning did not differ between the A-AB-A and A-B-A groups (t_(55)_ = -1.535, p = 0.302, CI for mean difference = [-10.883, 3.626]) at this time point. Likewise, there was no significant difference in direction error between these two groups at the end of learning either (t_(55)_ = -1.670, p = 0.302, CI for mean difference = [-7.501, 2.214], Figure 2e). Thus, reactivation of A did not increase interference during subsequent relearning.

We also examined whether there were any within group differences in A learning across days 1 and 3. A group x day ANOVA on the direction errors across all trials of A learning, revealed a significant interaction (F_(2,55)_ = 18.609, p < 0.001, ω^2^ = 0.092). Post-hoc tests confirmed that errors were significantly lower in the A-A group on day 3 than day 1 (t_(55)_ = 6.717, p < 0.001, CI for mean difference = [3.164, 8.487]). However, this was not the case for either the A-B-A group (t_(55)_ = - 1.692, p = 0.963, CI for mean difference = [-4.236, 1.225]) or the A-AB-A group (t_(55)_ = 0.640, p = 1.000, CI for mean difference = [-2.161, 3.300]), indicating that exposure to B eliminated the savings observed in the control group. Notably however, performance was not worse during relearning for either group.

Because reactivation did not increase interference, we next asked whether it might strengthen the original memory against interference. If so, participants in the A-AB-A group (who underwent reactivation) should have shown smaller direction errors on day 3 despite experiencing the interfering rotation B. However, this did not occur. One possibility is that the residual effects of B learning masked a potential benefit of reactivation during subsequent A relearning. To test this, these effects would need to be removed before A relearning on day 3. In a second experiment, we recruited two new groups: A-AB-NA (n = 18), who reactivated A before learning B, and A-B-NA (n = 19), who learned B without reactivation. In both groups, B was washed out with null trials before relearning A on day 3.

### Reactivation did not enhance relearning when competing learning was washed out after 24 hours

Figure 3a shows the learning curves for the two groups across the 3 days of the experiment. As in experiment 1, learning of A on day 1 across both groups was canonical and progressed as a gradual reduction in direction error. We found a main effect of time (F_(1,35)_ = 793.000, p < 0.001, ω^2^ = 0.862), no group (F_(1,35)_ = 0.438, p = 0.512, ω^2^ = 0) or interaction (F_(1,35)_ = 2.312, p = 0.137, ω^2^ = 0.010) effects in a group x time ANOVA (Figure 3b). When re-exposed to A on day 2, direction errors in the A-AB-NA group were significantly lower than naive A learning (Figure 3c), indicating that the previously acquired A memory had been retained and was being retrieved. This was supported by a day x time interaction (F_(1,17)_ = 84.297, p < 0.001, ω^2^ = 0.411), with post-hoc tests showing smaller direction errors during early learning (t_(17)_ = 10.240, p < 0.001, CI for mean difference =[8.494, 15.481]), but not later (t_(17)_ = 0.058, p = 0.954, CI for mean difference = [-1.668, 1.734]) when the naïve (day 1) and reactivation (day 2) periods were compared.

**Figure 3.**
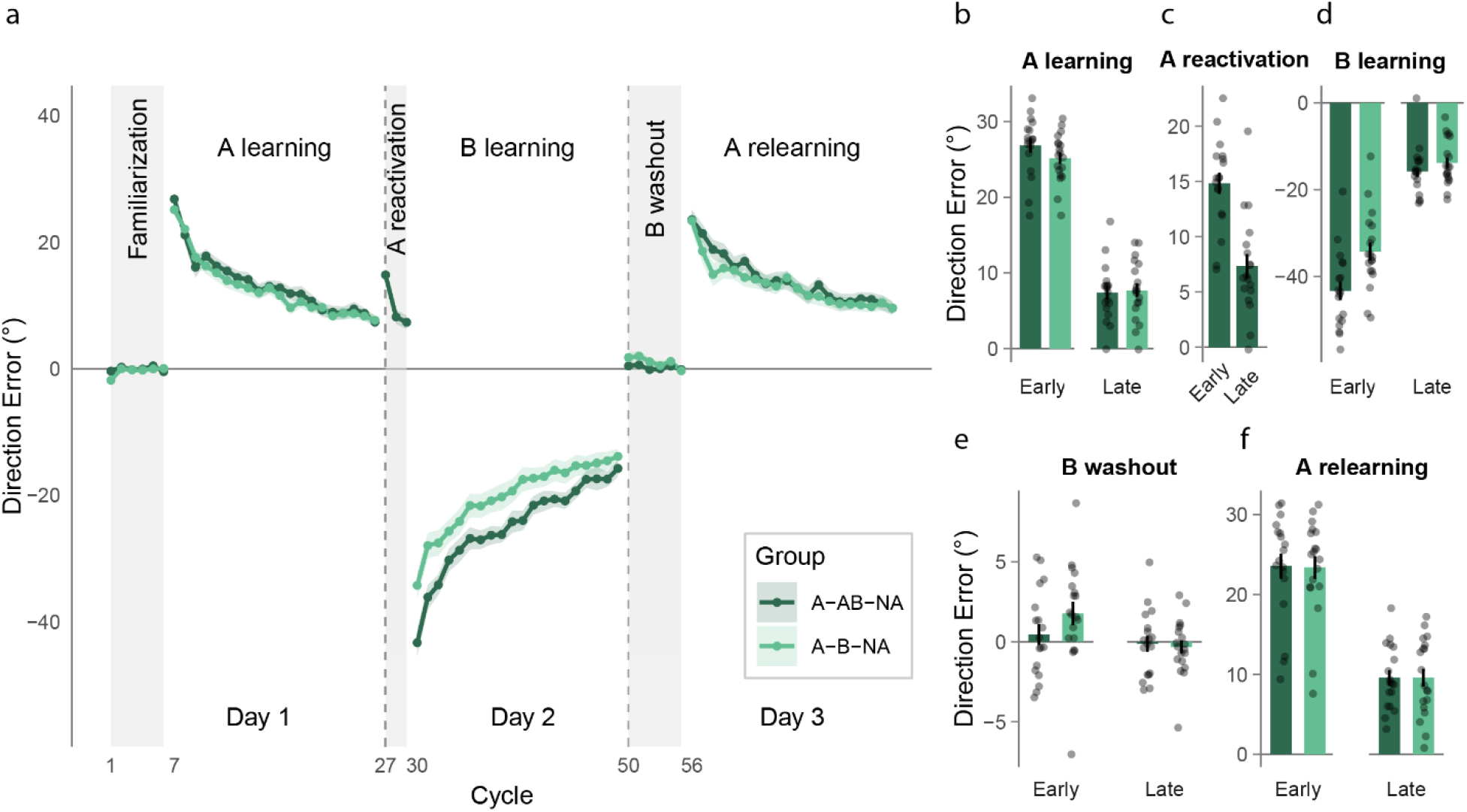
**(a)** Learning curves showing mean direction error as a function of cycles, plotted for the different blocks of experiment 2. The reactivation group (A-AB-NA) is shown in dark green, while the no-reactivation group (A-B-NA) is in light green. Dark vertical dashed lines represent 24-hour intervals between the blocks. Shaded areas represent SEM. **(b, c, d, e, f)** Mean ± SEM direction error during early and late phases of the different learning and washout blocks. Dots represent individual participants.

When exposed to B, both groups showed errors in the opposite direction with larger direction errors in the A-AB-NA group due to aftereffects of reactivating A (Figure 3d). We obtained a significant group x time interaction (F_(1,35)_ = 4.955, p = 0.033, ω^2^ = 0.046); post-hoc tests showed smaller direction errors in the A-B-NA compared to the A-AB-NA group during early B learning (t_(35)_ = -3.081, p = 0.008, CI for mean difference = [-17.825, -0.837]), but not at the end (t_(35)_ = - 1.072, p = 0.291, CI for mean difference = [-6.898, 3.075]). 24 hours after adapting to rotation B, null trials were introduced. These trials were introduced to remove potential anterograde effects of B. We found neither a group difference (F_(1,35)_ = 0.696, p = 0.410, ω^2^ = 0), nor a group x time interaction (F_(1,35)_ = 2.643, p = 0.113, ω^2^ = 0.015) in washout performance; only a significant time effect (F_(1,35)_ = 8.064, p = 0.007, ω^2^ = 0.060) was observed. This indicated that there was no difference in the washout pattern across groups (Figure 3e).

After washout, all participants readapted to A. We expected that if reactivation had protected the memory of A, it would be evident in the form of smaller errors on day 3 in the A-AB-NA group compared to the A-B-NA group. However, we found neither a main effect of group (F_(1,35)_ = 0.003, p = 0.955, ω^2^ = 0), nor a significant interaction (F_(1,35)_ = 0.007, p = 0.933, ω^2^ = 0) in a group x time ANOVA (Figure 3f). Only a significant effect of time was present (F_(1,35)_ = 203.669, p < 0.001, ω^2^ = 0.617). We also examined potential within group differences in A learning across days 1 and 3. Our group x day ANOVA, with data collapsed across all trials of A learning on days 1 and 3, revealed no significant effect of group (F_(1,35)_ = 0.317, p = 0.577, ω^2^ = 0), day (F_(1,35)_ = 3.308, p = 0.078, ω^2^ = 0.006) or group x day interaction (F_(1,35)_ = 0.052, p = 0.821, ω^2^ = 0). This indicated that relearning on day 3 in both groups was no different from their initial learning of A on day 1. Thus, reactivation did not improve relearning when B was washed out after a 24-hour delay.

We reasoned that delaying washout by 24 hours may have allowed the competing memory to stabilize, thereby obscuring any potential reactivation benefit. Therefore, in experiment 3, we washed out B immediately after learning in two new groups: A-ABN-A (with reactivation, n = 18) and A-BN-A (no reactivation, n = 18). Our main question was whether participants in the A-ABN-A group would demonstrate smaller errors compared to the A-BN-A group when relearning A on day 3, reflecting a reactivation-mediated benefit.

### Immediate washout of competing learning revealed a reactivation benefit

Figure 4a shows the learning curves for the two groups across the 3 days of the experiment. As observed in our first two experiments, adaptation to A on day 1 was typical; direction errors reduced from about 27° early on to about 9° by the end of the learning. There was a clear effect of time (F_(1,34)_ = 580.318, p < 0.001, ω^2^ = 0.832), but no group effect (F_(1,34)_ = 3.365, p = 0.075, ω^2^ = 0.033) or interaction (F_(1,34)_ = 0.378, p = 0.543, ω^2^ = 0) in a group x time ANOVA for day 1 A learning (Figure 4b). When exposed to A on day 2, direction errors in the A-ABN-A group were significantly lower than naïve A learning, as expected. This was supported by a significant day x time interaction (F_(1,17)_ = 175.535, p < 0.001, ω^2^ = 0.346). Post-hoc tests revealed smaller errors during early reactivation (t_(17)_ = 15.888, p < 0.001, CI for mean difference = [9.169, 13.409]) but not later (t_(17)_ = -0.507, p = 0.619, CI for mean difference = [-1.934, 1.372]) compared to naïve A learning of day 1 (Figure 4c).

**Figure 4.**
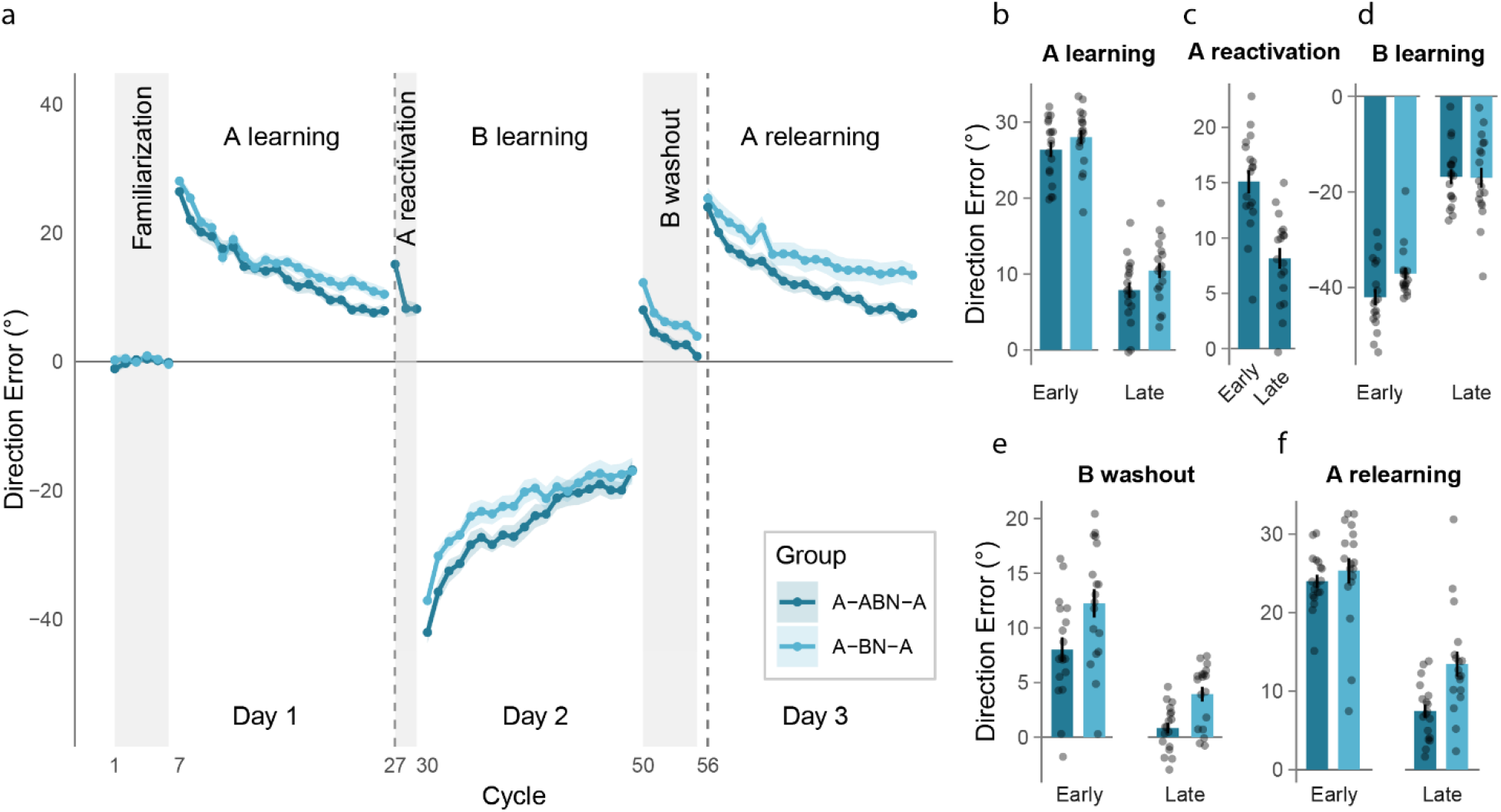
**(a)** Learning curves showing mean direction error as a function of cycles, plotted for the different blocks of experiment 3. The reactivation group (A-ABN-A) is shown in dark blue, while the no-reactivation group (A-BN-A) is in light blue. Dark vertical dashed lines represent 24-hour intervals between the blocks. Shaded areas represent SEM. **(b, c, d, e, f)** Mean ± SEM direction error during early and late phases of the different learning and washout blocks. Dots represent individual participants.

When exposed to B, both groups showed errors in the opposite direction, which decreased over trials. Although early direction errors in the A-ABN-A group (−42.049±1.669°) were greater than those in the A-BN-A group (−37.106±1.237°) reflecting aftereffects of the reactivated A memory (Figure 4d), there was no statistically reliable effect of group (F_(1,34)_ = 1.489, p = 0.231, ω^2^ = 0.007) or group x time interaction (F_(1,34)_ = 3.970, p = 0.054, ω^2^ = 0.025). There was only a main effect of time (F_(1,34)_ = 310.654, p < 0.001, ω^2^ = 0.728). This again indicated that participants in both groups adapted to B in a similar manner. B learning was then immediately washed out through null trials. During washout, we observed a significant effect of group (F_(1,34)_ = 10.825, p = 0.002, ω^2^ = 0.123) and time (F_(1,34)_ = 108.068, p < 0.001, ω^2^ = 0.485), but the group x time interaction was not significant (F_(1,34)_ = 0.574, p = 0.454, ω^2^ = 0). On average, direction errors in the A-BN-A group were greater by 3.672±1.116°, and this small difference was present at the beginning and at the end of the washout block. Critically, the two groups did not differ in terms of the magnitude of change in hand deviation that occurred over the washout block (from early to late) (t_(34)_ = -0.757, p = 0.454, CI for mean difference = [-4.157, 1.9]), indicating comparable removal of B-related adaptation before day-3 testing.

When participants relearned A on day 3, the group that had reactivated A on day 2 showed improved performance relative to the no-reactivation group. We noted significant group (F_(1,34)_ = 5.553, p = 0.024, ω^2^ = 0.061) as well as interaction (F_(1,34)_ = 6.452, p = 0.016, ω^2^ = 0.038) effects in a group x time ANOVA performed on the direction errors of A relearning. Post-hoc tests showed that there were no group differences at early learning (t_(34)_ = -0.752, p = 0.457, CI for mean difference = [-6.423, 3.704], Figure 4f); however, errors were significantly smaller for the A-ABN-A group during the late phase (t_(34)_ = -3.317, p = 0.004, CI for mean difference = [-11.050, -0.932], Figure 4f). We also probed within group differences in A learning across days 1 and 3. Our group x day analysis comparing the overall A learning between the two group across the two days, revealed a significant group effect (F_(1,34)_ = 5.115, p = 0.030, ω^2^ = 0.056) and a (marginal) interaction (F_(1,34)_ = 4.258, p = 0.047, ω^2^ = 0.013). Separate day-wise comparisons for each group showed that while direction errors during A learning were not different between days 1 and 3 in the A-BN-A group (t_(17)_ = -0.962, p = 0.349, CI for mean difference = [-3.115, 1.164]), they were significantly smaller on day 3 than day 1 for the A-ABN-A group (t_(17)_ = 2.558, p = 0.02, CI for mean difference = [0.246, 2.562]). Thus, multiple analyses converged to show that errors during relearning A on day 3 were smaller in the group that reactivated A on day 2.

As a final control, we tested whether performance during the reactivation phase differed across the groups. We performed a group x time analysis on the direction errors of the reactivation session of day 2. We found neither a significant group effect (F_(2,52)_ = 0.296, p = 0.745, ω^2^ = 0) nor a significant interaction (F_(2,52)_ = 0.118, p = 0.889, ω^2^ = 0). Only a significant effect of time (F_(2,52)_ = 215.684, p < 0.001, ω^2^ = 0.4) was evident, indicating that there was no difference in performance across the groups that underwent reactivation of A on day 2 (Figure 5). This ruled out the possibility that the advantage uncovered in experiment 3 was related to differences in the way reactivation proceeded across the groups.

**Figure 5.**
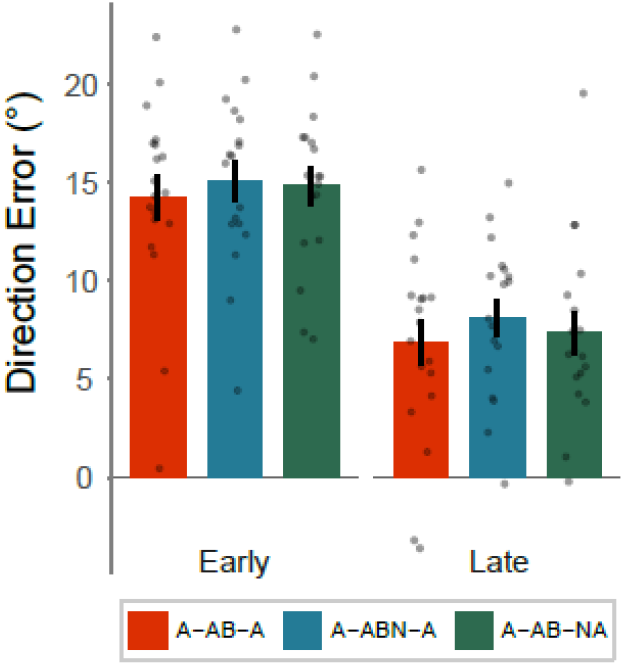
Mean ± SEM direction error during the early and late phases of the reactivation block for the three reactivation groups: A-AB-A (red), A-AB-NA (green) and A-ABN-A (blue). Dots represent individual participants.

## DISCUSSION

In this study, we examined how reactivation influences interactions between competing motor memories in error-based motor learning. Our findings suggest that interference between motor memories during adaptation does not depend on reactivation-induced destabilization of a stored memory. We obtained three main results. First, in line with past work (Caithness, 2004; Krakauer et al., 2005), learning opposing perturbations in succession produced substantial interference. Second, interference occurred regardless of whether the original memory had been explicitly reactivated, indicating that reactivation is not necessary for a competing memory to disrupt a previously acquired memory. Third, and most notably, reactivation *reduced* interference when acquisition of the competing memory was followed immediately by washout, which limited the opportunity for that memory to stabilize. Collectively, these findings suggest that interference reflects how competing motor memories are reinstated and expressed at retrieval, rather than a consequence of reactivation-induced destabilization.

The finding that interference is independent of reactivation is difficult to reconcile with a straightforward application of reconsolidation theory to motor adaptation. This framework proposes that active retrieval makes a previously consolidated memory temporarily labile and susceptible to modification by subsequent experience (Gabitov et al., 2017, 2019; Lee et al., 2017; Schiller & Phelps, 2011; Trent et al., 2015; Walker et al., 2003). Behaviorally, this would be expressed as poorer performance when the original memory is reassessed later. However, we found no evidence that reactivation amplified interference. Moreover, interference occurred even when the original A memory had not been explicitly reactivated. Taken together, these observations suggest that interference can occur without clear behavioral evidence for a reconsolidation-like destabilization process, and is therefore not a reliable indicator of destabilization. Notably, this aligns with some past studies of sequence learning that failed to observe reliable reconsolidation effects following brief memory reactivation (Censor et al., 2016; de Beukelaar et al., 2014; Hardwicke et al., 2016), raising questions about whether reactivation reliably renders motor memories labile.

An alternative to a destabilization-based account is that rather than transitioning between labile and consolidated states, motor memories transition between active and inactive states (Caithness, 2004; Censor et al., 2010; Lewis, 1979). In this view, previously acquired memories can be reinstated simply by re-engaging with the task, and this reinstatement brings the memory into an active state where it can be influenced by new learning. Thus, in our study, when participants returned to practice task B on day 2, the original memory (of A) may have returned to an active state in which it could interact with new learning (of B) even without its explicit retrieval or rehearsal. This aligns with broader theories of memory that emphasize contextual control over memory activation and subsequent expression across episodic, fear and sequence learning (Gershman et al., 2013; Hupbach et al., 2007; Nadel & Sederberg, 2024). However, this state-based account predicts similar susceptibility to interference regardless of whether A is explicitly reactivated or not. It therefore cannot readily explain our key result from experiment 3 that reactivation *reduces* interference when B is immediately washed out. This pattern suggests that reactivation does more than simply reinstate the prior memory. One possibility is that brief re-exposure to A incrementally *strengthens* the reactivated memory through additional practice. Thus, what is operationalized as “reactivation” may serve two separable functions: reinstating the prior memory and reinforcing it through further practice. This interpretation is consistent with evidence that even limited overtraining enhances resistance to interference (Krakauer et al., 2005; Shadmehr & Brashers-Krug, 1997). Yet, this account leaves open the important question of why the benefit of strengthening emerges only when washout of B is immediate.

This pattern cannot be explained by a simple timing-based account (e.g., differences in temporal spacing), because the interval between A and B learning was identical across experiments. One way to account for our findings, particularly the selective benefit observed in experiment 3, is via the Contextual Inference (COIN) model of motor learning (Heald et al., 2021). Here, learners infer latent motor contexts from the statistical structure of sensorimotor errors, and behavior reflects the weighted expression of these inferred contexts. Thus, reactivating A on day 2 may increase the prior probability assigned to the A context at the onset of B learning. This bias could promote separation of A and B into distinct latent states, reducing competition at retrieval. However, whether this separation is preserved depends on the subsequent fate of B. When B learning is washed out immediately after acquisition (experiment 3), its statistical structure is disrupted before it stabilizes, allowing the strengthened prior for A to remain dominant at retrieval. In contrast, when washout is delayed (experiments 1 and 2), B learning can form a stable contextual representation that competes with A at retrieval, masking any strengthening conferred by reactivation. Under this view, the protective effect of reactivation may emerge only when the competing memory is prevented from stabilizing into an independent contextual state. Memory expression may therefore reflect probabilistic contextual inference shaped by recent experience and consolidation dynamics, rather than changes in the stability of the underlying memory trace.

Alternative explanations are unlikely to account for the reactivation-mediated benefit observed in experiment 3. The broader task environment, apparatus, and movement structure were identical across groups, making simple contextual differences unlikely to explain the effect. Moreover, the initial conditions on day 2 were identical in experiments 2 and 3, yet reactivation-mediated benefits emerged only when washout occurred immediately after B learning. Finally, both groups exhibited comparable error levels at the start of A relearning on day 3 (approximately 25° and 24°), with the difference between groups emerging only over the course of relearning. This pattern suggests that the benefit of reactivation was not present at the outset, but emerged progressively over the course of subsequent adaptation.

It is not uncommon in reconsolidation studies across various memory domains to draw a hard distinction between reactivation-mediated protection of a prior memory or its weakening (Forcato et al., 2010, 2011; Levy et al., 2018; Sinclair & Barense, 2019). This likely reflects the fact that the experimental paradigms that probe strengthening and weakening are often fundamentally different. Rarely do the studies demonstrating protection or strengthening challenge the reactivated memory with a competing one, and seldom does a behavioral paradigm demonstrating weakening of an existing memory trace not use a competing input. Our results suggest that this binary framework is overly simplistic; we show that “strengthening” can manifest as greater resistance to interference from a competing input. In other words, the protective effect of reactivation does not merely involve enhancing memory stability in isolation but may also involve shielding the memory from being disrupted by new, conflicting information. This suggests that the traditional dichotomy between strengthening and weakening may be too restrictive. Instead, reactivation may serve a more nuanced role, at times stabilizing memories and increasing their resistance to interference.

Our results also raise several questions about the conditions under which reactivation strengthens motor memories. In animal studies, both weakening and strengthening effects are known to be governed by a set of “boundary conditions”, which include, among others, the age of the memory, the duration of reactivation, and the novelty of information introduced during reactivation (de Beukelaar et al., 2014, 2016; Fernández et al., 2016; Mooney et al., 2021; Nadel & Sederberg, 2024; Ritvo et al., 2019; Sinclair & Barense, 2019). Whether similar constraints govern reactivation effects in human motor learning remains unclear and remains to be tested. Future studies can also probe how reactivation interacts with the distinct explicit and implicit components of motor learning (McDougle et al., 2015; Oza et al., 2024). Finally, manipulations such as increased sensorimotor variability during reactivation have been shown to strengthen motor memories (Wymbs et al., 2016) and may provide a useful tool for probing the mechanisms underlying reactivation-mediated resistance to interference. Understanding these conditions may ultimately inform the development of more effective protocols in skill learning and rehabilitation, where minimizing interference from competing inputs and enhancing retention remains a central goal.

## Acknowledgements

We thank Devangshu Nandi, Hannah Jaison and Suryakant Kad for assistance with data collection, and Gaurav Panthi and Ajay Sahu for help with figure preparation. This work was supported by grants from the Department of Science and Technology, Government of India. Support from IIT Gandhinagar is also acknowledged.

